# Long-Term Potentiation Produces a Sustained Expansion of Synaptic Information Storage Capacity in Adult Rat Hippocampus

**DOI:** 10.1101/2024.01.12.574766

**Authors:** Mohammad Samavat, Thomas M. Bartol, Cailey Bromer, Dusten D. Hubbard, Dakota C. Hanka, Masaaki Kuwajima, John M. Mendenhall, Patrick H. Parker, Jared B. Bowden, Wickliffe C. Abraham, Terrence J. Sejnowski, Kristen M. Harris

## Abstract

Long-term potentiation (LTP) has become a standard model for investigating synaptic mechanisms of learning and memory. Increasingly, it is of interest to understand how LTP affects the synaptic information storage capacity of the targeted population of synapses. Here, structural synaptic plasticity during LTP was explored using three-dimensional reconstruction from serial section electron microscopy. Storage capacity was assessed by applying a new analytical approach, Shannon information theory, to delineate the number of functionally distinguishable synaptic strengths. LTP was induced by delta-burst stimulation of perforant pathway inputs to the middle molecular layer of hippocampal dentate granule cells in adult rats. Spine head volumes were measured as predictors of synaptic strength and compared between LTP and control hemispheres at 30 min and 2 hr after the induction of LTP. Synapses from the same axon onto the same dendrite were used to determine the precision of synaptic plasticity based on the similarity of their physical dimensions. Shannon entropy was measured by exploiting the frequency of spine heads in functionally distinguishable sizes to assess the degree to which LTP altered the number of bits of information storage. Outcomes from these analyses reveal that LTP expanded storage capacity; the distribution of spine head volumes was increased from 2 bits in controls to 3 bits at 30 min and 2.7 bits at 2 hr after the induction of LTP. Furthermore, the distribution of spine head volumes was more uniform across the increased number of functionally distinguishable sizes following LTP, thus achieving more efficient use of coding space across the population of synapses.

**Significance:** Establishing relationships between structure, function, and information storage capacity provides a new approach to assessing network strength from structural measurements. Long term potentiation (LTP) is a standard model for investigating synaptic mechanisms of learning and memory. Information is a retrievable quantity that is being stored in synapses as synaptic strength and is correlated with multiple structural components of synaptic strength. Structural synaptic plasticity was measured in 3D reconstructions from serial section electron microscopy of spine head volume, as a proxy for synapse strength, at 30 min and 2 hr after LTP induction. Outcomes indicate that LTP enhances information storage capacity for at least 2 hr by increasing the precision of the synaptic structure and expanding the range of synapse sizes.

## Introduction

Long-term potentiation (LTP) is widely studied to understand cellular, synaptic, and molecular mechanisms of learning and memory. Connecting synapse structure to function is a central question. Prior studies have demonstrated multiple changes in the structure of dendritic spines and synapses following the induction of LTP in the hippocampal dentate gyrus (Fifková and Van Harreveld, 1977; Fifková and Anderson, 1981; Desmond and Levy, 1986; Stewart et al., 2000; Geinisman et al., 1991; Weeks et al., 1999, 2000, 2001; Mezey et al., 2004; Trommald et al., 1996; Popov et al., 2004). The structural effects emerged as early as 30 min, and some changes lasted for hours to days following induction of LTP (Weeks et al., 2001; Popov et al., 2004; Medvedev et al., 2010). Although structural synaptic plasticity is well-established as an experience-dependent mechanism for modifying synaptic features, the precision of this mechanism and its effects on the underlying capacity for information storage are unknown.

From an information theory perspective, there can be no information stored without precision. To determine precision, differences in the structure of synapses that have the same history of activation are measured because their structures are predicted to be similar. Synaptic strength is highly correlated with dendritic spine head volume (Harvey and Svoboda, 2007; Matsuzaki et al., 2004; Harris, 2020; Yang and Lui, 2022). Thus, the dimensions of pairs of dendritic spines on the same dendrite that receive input from the same axon are expected to be similar due to similar activation histories (Sorra and Harris, 1993; Koester and Johnston, 2005; Trommald et al., 1996; Popov and Stewart, 2009; Bartol et al., 2015, Kumar et al., 2020; Kasthuri et al., 2015; Dvorkin and Ziv, 2016; Bloss et al., 2018; Motta et al., 2019; Dorkenwald et al., 2019). Differences in the dimensions of these same-dendrite same-axon (SDSA) pairs will reflect natural or transient variation in synapse structure that is not due to differences in overall functional status. This variation can then be used to provide boundaries around spine dimensions that have essentially the same efficacy. Signal detection theory was used in our previous studies to estimate the number of distinguishable synaptic strengths based on this spine head volume measure of precision (Bartol et al., 2015; Bromer et al., 2018). Here, a more robust approach, based on Shannon information theory, was used to overcome the limitations of signal detection theory (Samavat et al., 2023). The findings provide new evidence that the structural changes associated with LTP last at least 2 hours and enhance information storage capacity and efficiency.

## Results

### LTP and Control datasets

The delta-burst protocol was used to produce LTP on the medial perforant pathway in the middle molecular layer of the dentate gyrus, and control responses were measured in the contralateral hippocampus from the same animals (Bowden et al., 2012). Relative to their baseline responses, the LTP hemispheres showed an average of 41% potentiation in the 30 min experiments (Fig. 1A, B) and 37% potentiation for the 2 hr experiments (Fig. 1C). In both the 30 min and 2 hr experiments, responses in the control hemispheres were stable across time. Representative serial section electron micrographs illustrate excellent tissue preservation for all conditions, and 3D reconstructions of representative dendrites show the full segment reconstructions surrounding the central regions that were analyzed for the SDSA pairs (Fig. 1D-G). Spine and axon densities were comparable for the 30 min and 2 hr control and LTP datasets (Table 1). The density of SDSA pairs was comparable across 30 min and 2 hr controls, but somewhat lower in the 30 min and 2hr LTP conditions (Table 1). This shift may reflect an LTP effect, however, given the small sample size it may also reflect variation among dendritic samples not related to LTP (Table 1). These datasets proved sufficient to detect LTP related changes in spine head volumes, SDSA precision, and synaptic information storage capacity.

**Figure 1:**
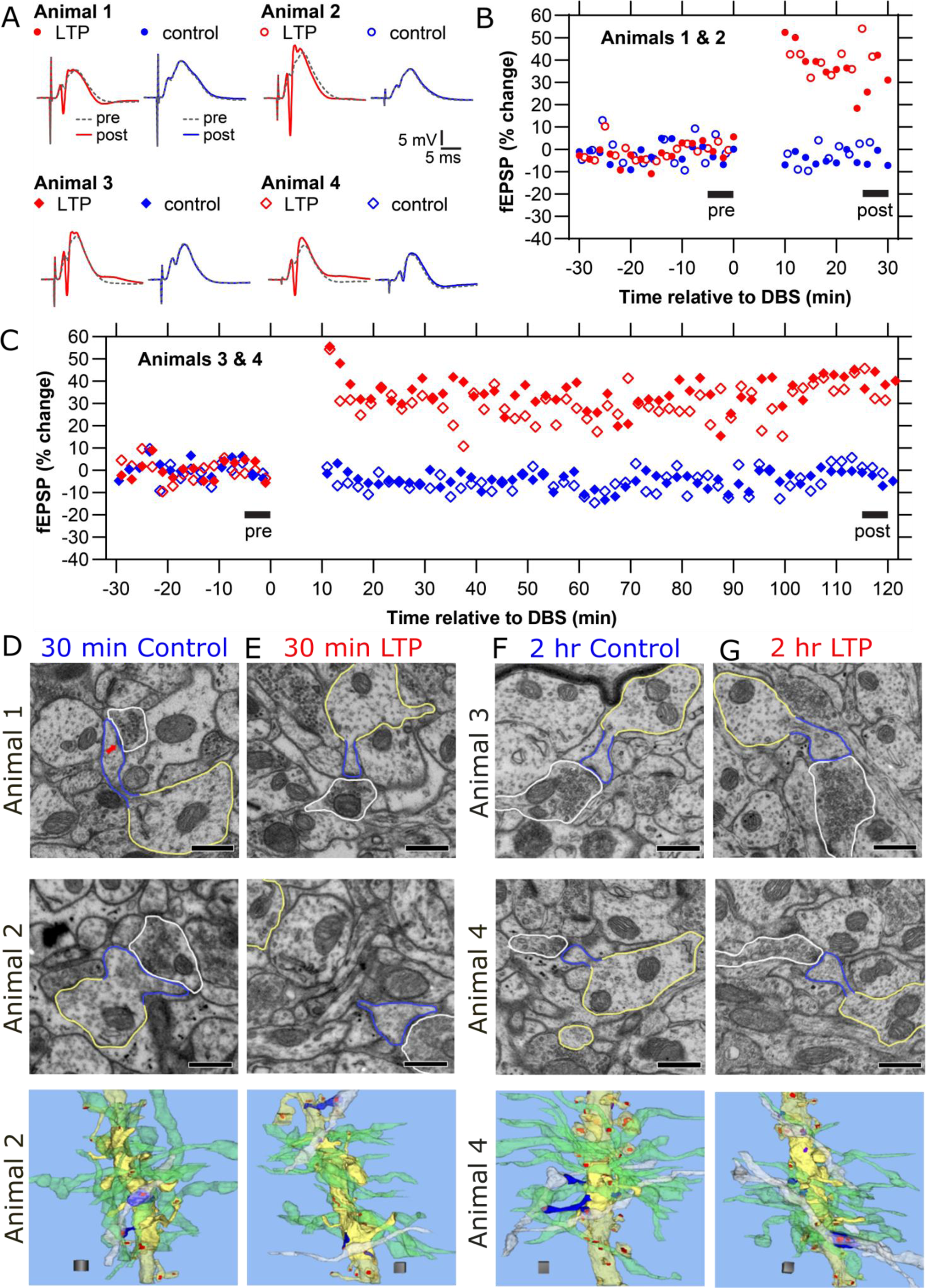
LTP and control responses and sample dendrites from each condition. (A) Representative waveforms from baseline responses (dotted, pre) superimposed on responses following delta-burst stimulation (solid, post) in the LTP (red) or control (blue) hemispheres. (B) The fEPSP responses during the 30 min experiment shows percentage change after induction of LTP (red) at time 0, and stable control responses (blue). (C) The fEPSP responses during the 2 hr experiment show percentage change after induction of LTP (red) at time 0, and stable control responses (blue). (D-G top two rows) Example electron micrographs from each animal and condition, illustrating dendritic shaft (yellow), dendritic spine (blue), axon (white), and PSD (red arrow). (Scale bars = 0.5 µm.) (D-G bottom row) 3D reconstructions of dendrites and spines (yellow) with excitatory (red) and inhibitory (purple) synapses. Most of the axons (green) synapsing on the 15 spines along the middle of the dendrite (solid yellow) made synapses with just one spine, while one or two axons (white) made synapses with two spines (blue). The dendrite and spines occurring along the rest of the reconstructed segment are illustrated in translucent yellow. (Scale cube = 1 µm^3^.) Videos are provided for D-G.

**Table 1:** Datasets by condition.

| Condition | Total Length (μm) | Analyzed Length (μm) | #Spines | #Axons | #SDSA pairs | #Spines/μm | #Axons/μm | #SDSA Pairs/μm |
| --- | --- | --- | --- | --- | --- | --- | --- | --- |
| 30 min Con | 8.6 - 10.6 | 3.4 - 6.7 | 13 - 19 | 14 - 17 | 1 - 5 | 2.3 - 4.4 | 2.1 - 4.4 | 0.2 - 1.2 |
| 30 min LTP | 9.3 - 10.6 | 3.7 - 6.8 | 13 - 19 | 16 - 20 | 1 - 2 | 2.2 - 4.45 | 2.5 - 4.7 | 0.2 - 0.5 |
| 2 hr Con | 8.0 - 10.9 | 2.7 - 8.4 | 12 - 19 | 15 - 24 | 2 - 5 | 2.0 - 4.5 | 2.4 - 5.5 | 0.2 - 1.5 |
| 2 hr LTP | 8.6 - 11.3 | 4.7 - 6.4 | 14 - 17 | 15 - 18 | 0 - 3 | 2.6 - 3.1 | 2.3 - 3.9 | 0 - 0.6 |

All data are expressed as the range from minimum to maximum values. Columns include total length (fully reconstructed dendritic segment length); analyzed length (length of the analyzed dendritic segment for determining SDSA pairs); number of spines, axons, and SDSA pairs along the analyzed length; number per micron frequency of spines, axons, and SDSA pairs along the analyzed length.

### Impact of LTP on spine head volumes

Small and large dendritic spines had diverse shapes and a broad range in spine head volumes (Fig. 2, 3A-D). Differences between the LTP and control histograms revealed significant increases in small and large spine frequencies for LTP relative to control stimulation at both 30 min and 2 hr following the induction of LTP (Fig. 3E, Table 2). By 2 hr the peaks and troughs shifted relative to 30 min such that the increase in smaller spines was significantly reduced and the increase in larger spines was stabilized during LTP (Fig. 3F, Table 2).

**Figure 2:**
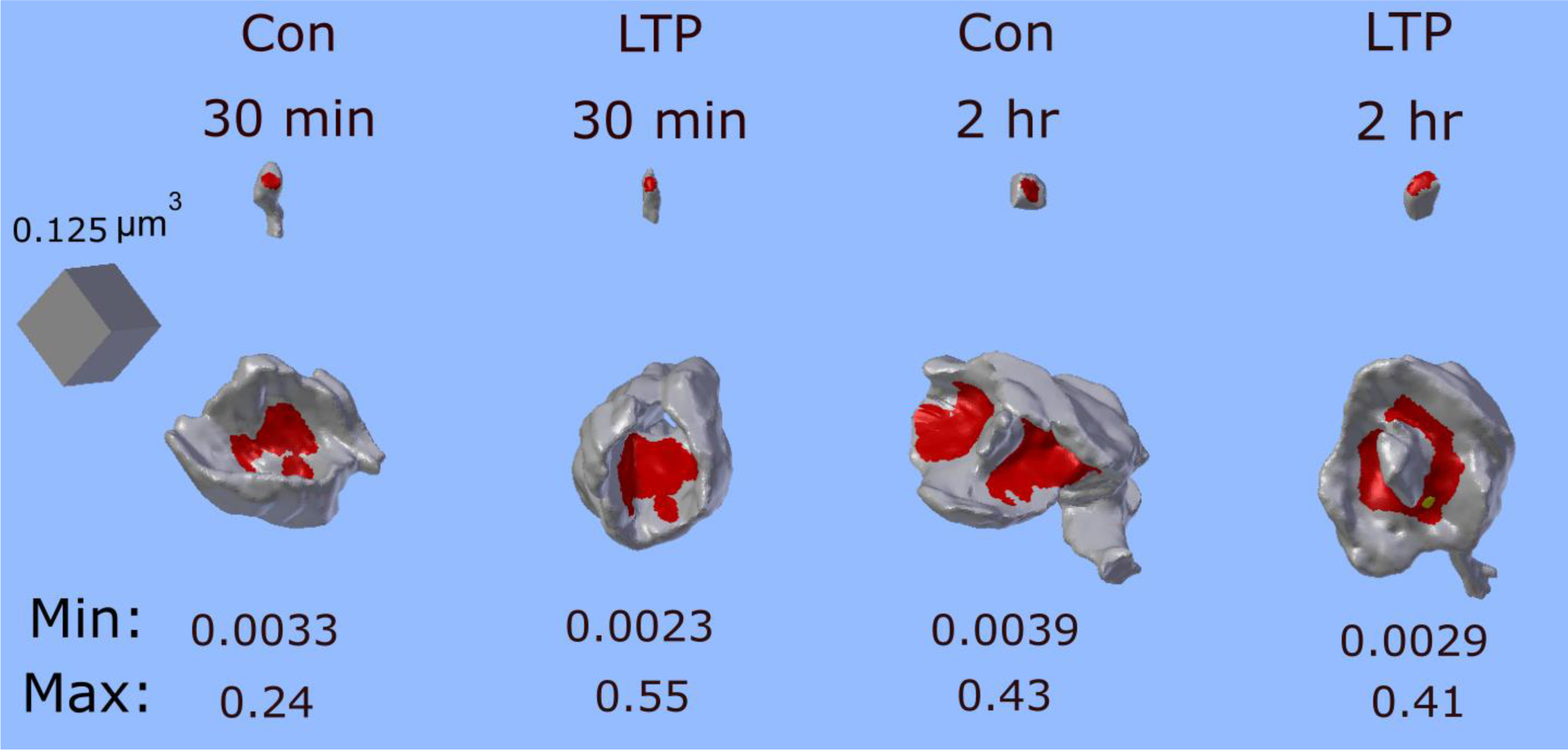
Range in spine head volumes from each condition based on 3D reconstruction of the smallest and largest spine redheads (spine head gray, synapse) in each condition. (Minimum and maximum volumes indicated in *μ*m^3^.)

**Figure 3:**
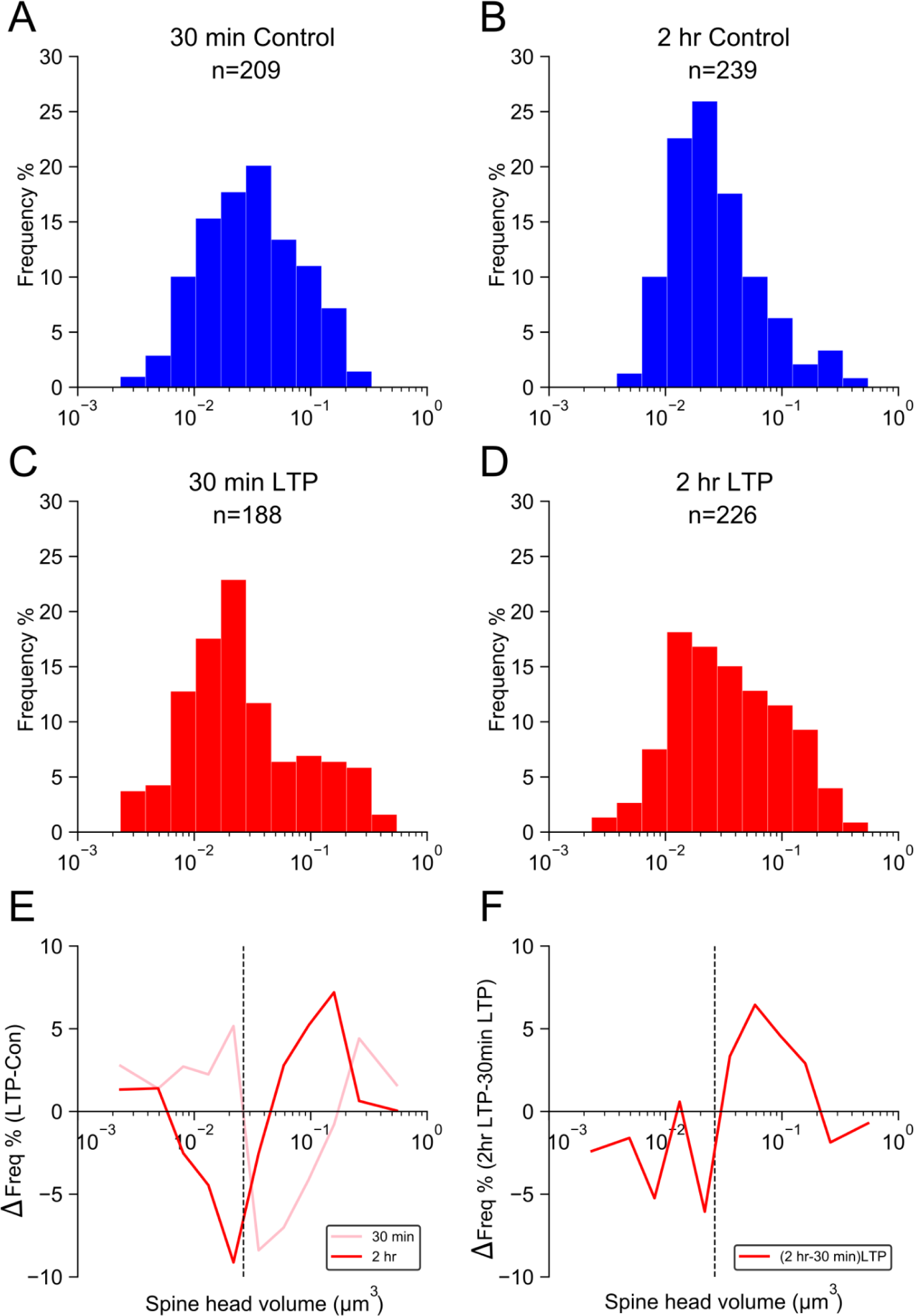
Change relative to control hemispheres in the distribution of spine head volumes at 30 min and 2 hr after the induction of LTP. (A-D) Frequency distributions of spine head volumes from control and LTP hemispheres divided across 11 equal sized bins on a log scale, since the raw measures are highly skewed. (E) Difference between the frequency of spine head volumes in control and LTP conditions (i.e., LTP - control) at 30 min (pink) and 2 hr (red). (F) Difference between the frequency of spine head volumes in 30 min LTP and 2 hr LTP conditions. The vertical dashed line is the median of combined spine head volumes. All peaks and troughs in E and F are statistically significantly different from 0% change (P < 0.005).

**Table 2:** P values of the observed peaks and troughs in Fig 2 E,F relative to 10,000 bootstrap values.

| Test Statistic | SHV<median | P-Value | SHV>median | P-Value |
| --- | --- | --- | --- | --- |
| sum (2 hr LTP - 30 min LTP) | -15.6% | 0.0015 | 16.1% | 0.0006 |
| sum (2 hr LTP- 2 hr Con) - (30 min LTP - 30 min Con) | -28.6% | 0 | 29.1% | 0 |
| sum (2 hr LTP - 2 hr Con) | -13.1% | 0.0025 | 13.2% | 0.0021 |
| sum (30 min LTP - 30 min Con) | 15.5% | 0.0011 | 16.0% | 0.0008 |

These histograms reflect a traditional way for describing changes in the relative frequencies between datasets. However, they do not reveal whether the shifts in spine size represent significant shifts in distinguishable functional status and thus storage capacity. To overcome this limitation, we applied new methods based on Shannon information theory (Samavat et al., 2023).

### Effect of LTP on Synaptic Information Storage Capacity

The concept of entropy comes from the field of thermodynamics and measures the number of distinguishable configurations of a system. To determine the number of functionally distinguishable synapse sizes, we first had to determine the precision of synaptic strengths. A single axon synapsing with 2 spines on the same short dendritic segment is the most likely arrangement to have synapse dimensions that reflect synchronous activation histories and thus similar strengths. The precision of spine head volumes was determined by calculating the median CV and standard error of the SDSA spine pairs using bootstrapping for each condition (Fig. 4). There was no significant trend in the value of the CV from the smallest to the largest spine head volumes (Fig. 4). Thus, synaptic plasticity based on co-activation history was as precise among small spines as it was among large spines for all control and LTP datasets.

**Figure 4:**
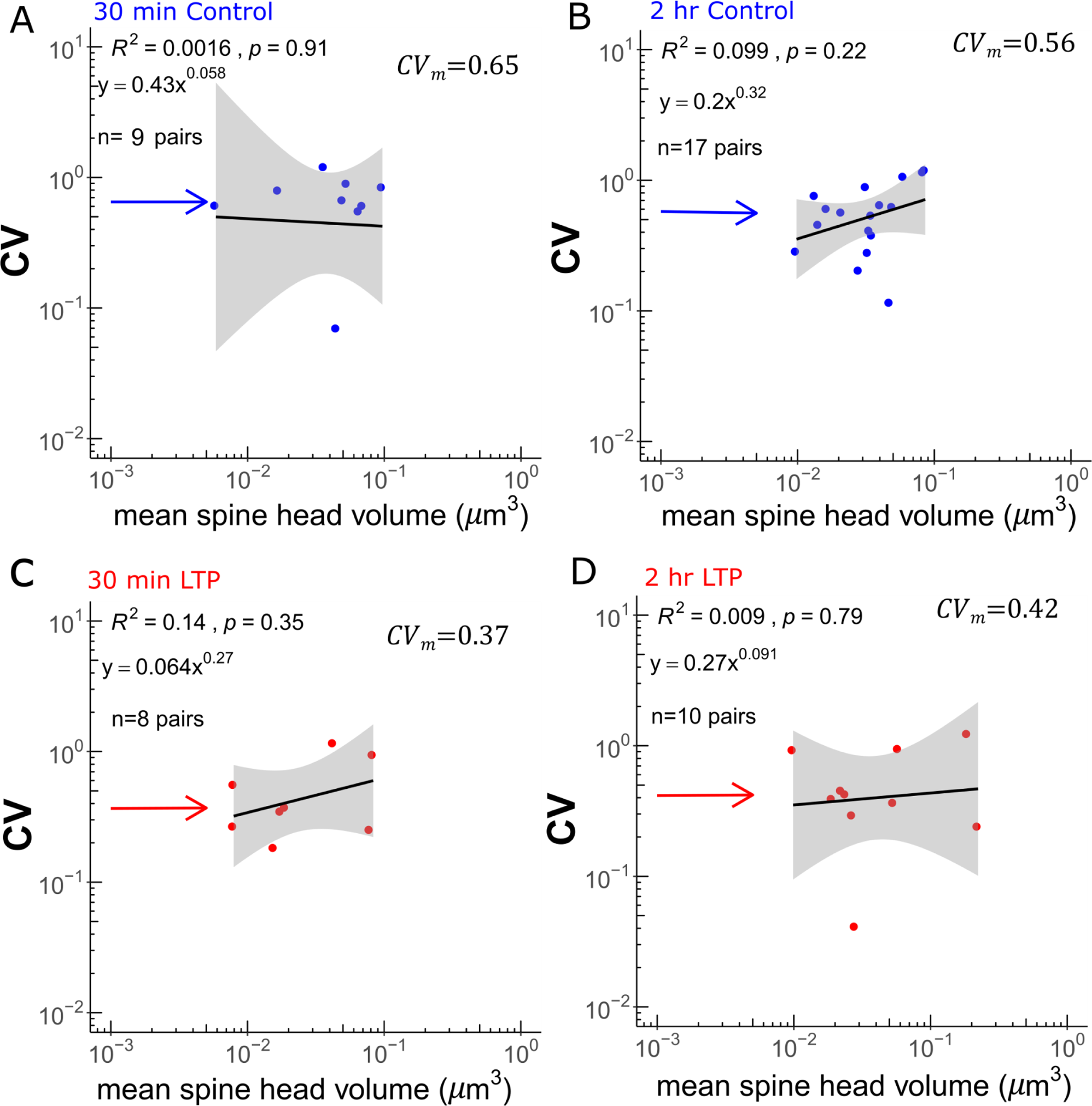
Analysis of synaptic precision based on the CV of SDSA pairs across time and plasticity. (A, B) 30 min and 2 hr control conditions. (C, D) 30 min and 2 hr LTP conditions. The gray region is the 95% confidence interval for each regression line. The Y axis is the CV for each SDSA pair. Arrows indicate the median CV (CV_m_) of the SDSA pairs for each condition. The X axis shows the mean value of the spine head volumes, on a log scale, for each SDSA pair. The one factor KW test was used to test for differences between the control and LTP CVs and showed no significant difference between the four conditions (*p*=0.16). The lowest CV value in the 30 min control (0.040) and 2 hr control (0.078) were treated as outliers and removed from subsequent analyses. (See Methods section for details of the bootstrap analysis.) It is worth noting that the average measurement error of the same spine head volumes among 4 investigators had a CV <0.01, which is much less than the median CV of these SDSA pairs.

The boundaries of each bin holding functionally distinguishable spine sizes were determined by the median CV of the SDSA pairs for each condition. The values contained in the first bin included the smallest spine head volume and all measured spines of greater size within one CV. Subsequent bins repeated this process to include sequentially larger spines within the CV of each bin (Methods). The last bin contained fewer spines than a full CV because it ended with the largest spine head volume in each condition (Fig. 5). The distribution of functionally distinguishable sizes is different from the histogram of spine head volumes (Fig. 3) because each bin is determined by the median CV for each condition, rather than a set bin width. The 30 min control had 5 functionally distinguishable sizes, which was comparable to 6 distinguishable sizes in the 2 hr control and demonstrated the similarity across rats and time-points (Fig. 5A, B). By 30 min after LTP induction, the number expanded to 10 distinguishable sizes (Fig. 5C). By 2 hr after LTP induction, the expansion remained elevated, but was reduced to 8 distinguishable sizes (Fig. 5D). At both of the 30 min and 2 hr time points, the elevation in distinguishable sizes due to LTP resulted from both an expansion in the range of spine sizes and a decrease in the CV of the SDSA pairs relative to their respective control values (Table 3).

**Figure 5:**
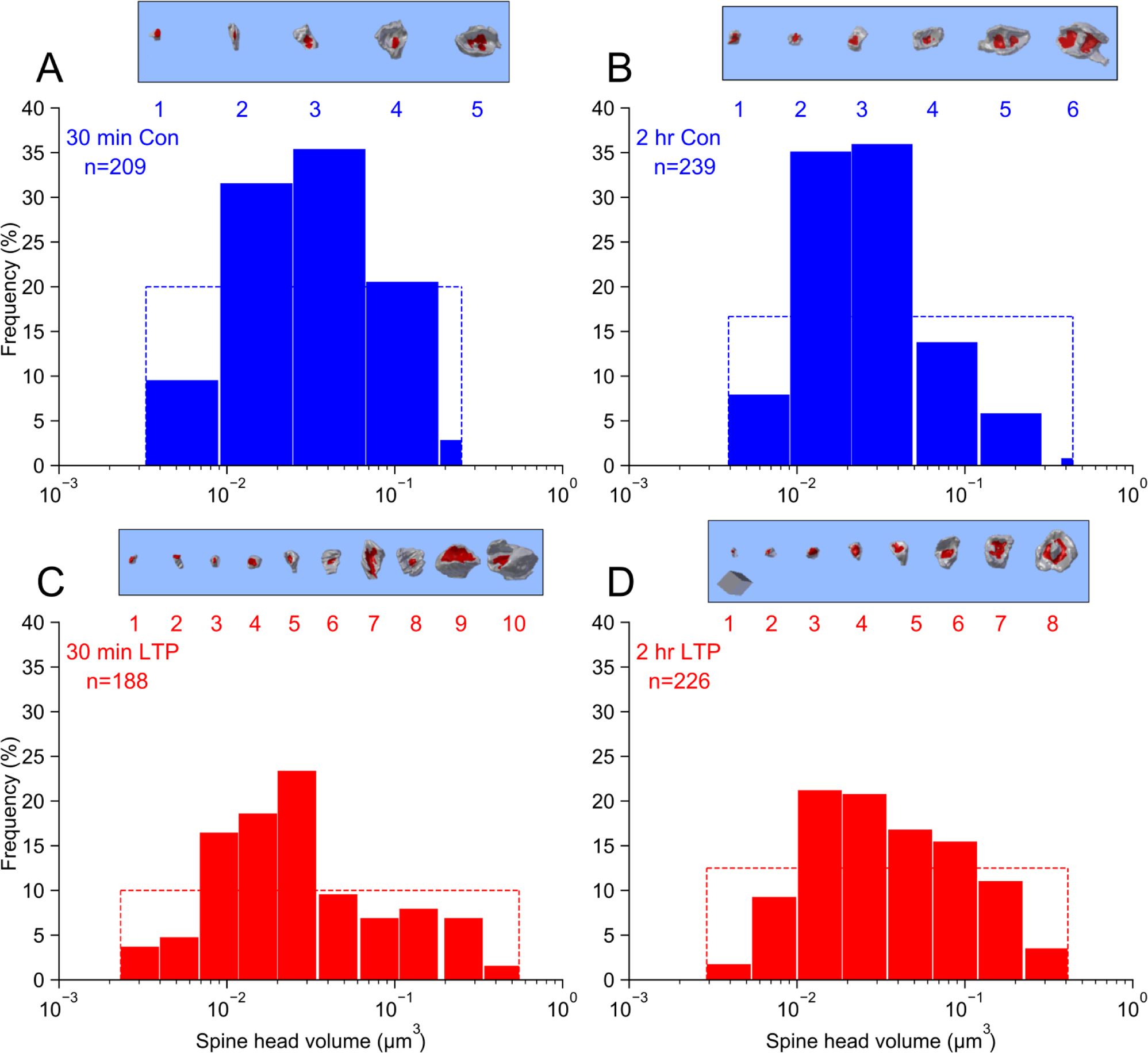
Distribution of distinguishable bins. (A-D) The X axis equals spine head volumes on a log scale. The widths of the bars represent distinguishable bins and are equal to [x1,x2], CV(x1,x2)= CV_m_ of the dataset, where x1 is the smallest spine head volume and x2 is the next larger hypothetical head volume that has the median CV with x1, except for the last state being truncated at the largest spine head volume. The Y axes indicate the percentages of spine head volumes in each state relative to the total number of spines in each dataset. The insets above each graph show the largest spine head in each state (cube in D for A-D = 0.125 µm^3^.) The dashed line indicates the theoretical uniform distribution with maximum entropy for each dataset. The sum of frequencies under each uniform distribution equals 100%.

**Table 3:** The number of distinguishable synaptic strength (N_s_) based on spine head volumes.

| Condition | # SDSA pairs | Median CV | # SHV | SHV ($\mu\text{m}^3$) | $N_s$ |
| --- | --- | --- | --- | --- | --- |
| 30min Con | 10 | $0.65 \pm 0.12$ | 209 | $0.031 \pm 0.0037$ | $5 \pm 0.24$ |
| 30min LTP | 8 | $0.37 \pm 0.16$ | 188 | $0.022 \pm 0.0014$ | $10 \pm 0.51$ |
| 2hr Con | 18 | $0.56 \pm 0.09$ | 239 | $0.023 \pm 0.0013$ | $6 \pm 0.48$ |
| 2hr LTP | 10 | $0.42 \pm 0.15$ | 226 | $0.031 \pm 0.0036$ | $8 \pm 0.20$ |

All data are expressed as median. For column 3 and 5 data are median ± standard error of the median, determined by calculating the standard deviation of the 1000 estimated median values of the 1000 bootstrap samples from empirical values with the same length. The standard error on the # of distinguishable synaptic strength ( $N_s$ ) was estimated by applying the clustering approach to each of the 1000 bootstrap samples (each sampled with replacement with the same size from empirical spine head volumes).

The amount of Shannon entropy was computed using the probability of occurrence of each size divided by the total number of measured spine head volumes. The Shannon entropy equaled 2 bits for the control datasets and increased to 3 and 2.74 bits for the LTP datasets at 30 min and 2 hr, respectively (Table 4). Maximum entropy is achieved when every distinguishable size has the same frequency as every other distinguishable size. The maximum entropy was calculated as log2(N_S_) where N_S_ is the number of distinguishable sizes. The distinguishable sizes of spine head volumes were then compared to this uniform distribution with maximum entropy, illustrated as dotted rectangles for each condition (Fig. 5). The Kullback-Leibler (KL) divergence statistic was used to assess the degree of divergence from maximum entropy. The KL divergence from uniform was about equal for the 30 min control and LTP datasets. In contrast, the KL divergence for the 2 hr LTP condition was less than half of its matched control condition (Table 4 overall, and Supplementary Table 1 by animal and condition).

**Table 4:** Calculating Shannon entropy in the frequency of distinguishable synaptic strength based on spine head volumes.

| Dataset | mean SHVs | CV SHVs | Shannon Entropy bits<br>$H(P) \pm \text{SE}$ | Maximum Entropy<br>$H(U) \pm \text{SE}$ | KL (P\|\|U)<br>($\pm \text{SE}$) | Efficiency Ratio $\eta$ |
| --- | --- | --- | --- | --- | --- | --- |
| 30min Con | $0.049 \pm 0.0037$ | $1.24 \pm 0.077$ | $2.0 \pm 0.32$ | $2.32 \pm 0.34$ | $0.33 \pm 0.1$ | 86% |
| 30min LTP | $0.053 \pm 0.0067$ | $1.72 \pm 0.15$ | $3.0 \pm 0.42$ | $3.32 \pm 0.42$ | $0.32 \pm 0.089$ | 90% |
| 2hr Con | $0.040 \pm 0.0034$ | $1.084 \pm 0.081$ | $2.05 \pm 0.27$ | $2.59 \pm 0.29$ | $0.53 \pm 0.10$ | 79% |
| 2hr LTP | $0.060 \pm 0.0052$ | $1.41 \pm 0.089$ | $2.74 \pm 0.41$ | $3.0 \pm 0.41$ | $0.26 \pm 0.086$ | 91% |

For columns 2-6 the data are presented as median ± standard error. The standard error is calculated by concurrently applying the estimator at the top of each column to each of the 1000 bootstrap samples (sampled with replacement from the empirical data). Then the standard deviation of the resulting 1000 estimated values reported as the standard error of the estimate from the empirical data. Note that $\text{KL}(P||U)/H(U)$ is equivalent to $\text{KL}(P||U)/\text{KL}_{\text{MAX}}$ .

The efficiency of the distinguishable sizes in storing information across a population of synapses was determined by the ratio of the measured KL divergence to its hypothetical minimum, i.e., maximal entropy (see Statistical Methods). Relative to within-animal controls, the efficiency ratio was increased 4% at 30 min and 12% at 2 hr, after the induction of LTP (Tables 4 and 5). Thus, spine synapses were not on/off 1-bit switches, and LTP increased the distinguishable synaptic sizes based on spine head volume towards the maximum information storage capacity with high efficiency.

## Discussion

In neuroscience, information theory has long been applied to the analysis of spike trains (Dayan and Abbott, 2005). This is the first report using Shannon information theory to assess information storage capacity of synapses based on structure and its change after the induction of LTP. At 30 min following the induction of LTP, spine head volumes shifted from the middle of the range towards both smaller and larger sizes. By 2 hr after LTP induction, the frequency of large spine head volumes was further elevated. The KL divergence from a uniform distribution was much less after LTP relative to controls at 30 min and was further reduced at 2 hr after LTP induction. Thus, the shifts produced a more uniform distribution of spine head volumes and increased Shannon entropy and coding efficiency. The Shannon information content increased from 2.0 bits in both sets of control synapses to 3.0 bits by 30 min after the induction of LTP and was sustained at 2.7 bits for at least 2 hr, indicating a sustained increase in synaptic information storage capacity.

Information storage capacity depends on the precision of distinguishable sizes that a synapse could assume (Samavat et al., 2023). In this study we used spine head volume as a proxy for synaptic strength because it correlates with other measures of synaptic strength, such as postsynaptic potentials, PSD area, and vesicle number, and is reliably measured across experimenters (Bartol et al., 2015). To distinguish each size, the precision is determined by measuring how similar two synapses are in size that have the same function. This precision was estimated by the covariance of spine head volumes in SDSA pairs because these pairs are the most likely to have had the same activation histories (Bartol et al., 2015; Bromer et al., 2018; Samavat et al., 2023). Importantly, there was no significant difference in median CV from pair to pair across the full range of spine sizes. There was also no significant difference in the median CV between LTP and control SDSA pairs, suggesting that as one spine of the pair enlarged (or shrinks), so did the other. Nevertheless, LTP produced a subtle decrease in CV, hence we used the median values from each dataset to set the boundaries around each distinguishable size.

Shannon information theory provides several advantages over signal detection theory, which was previously used to investigate synaptic information storage capacity (Bartol et. al, 2015; Bromer et al., 2018; Samavat et al., 2023). First, signal detection theory assumes that the employed Gaussian curves are distributed equally along the full range of spine head volumes without accounting for gaps in the distribution, and thus tends to overestimate the true number of distinguishable synaptic sizes. The Shannon information theory approach excludes gaps and converges toward the true number of distinguishable sizes as more of the distribution is sampled. Indeed, the number of bits of information in the 30 min datasets in the current study is smaller than reported by Bromer et al. (2018), although the effect of LTP induction on this measure is similar. A second advantage is that the full population of spine head volumes are included in the distribution, greatly improving the statistical power of the estimate over the signal detection approach where only the spines involved in the SDSA pairs defined the range (Bartol et al., 2015). A third advantage of our new approach is that there are no free parameters in the estimate, unlike signal detection theory where the degree of overlap of the Gaussians is a free parameter. A fourth advantage is that the new method is robust to outliers of skewed distributions, adding only one distinguishable size per outlier, unlike equally spaced Gaussians that assign bits to nonexistent intervening values. Finally, this new approach reveals the frequency of spines in each distinguishable size, allowing calculation of modes, gaps, entropy, and efficiency of distribution across distinguishable synaptic strengths.

An important constraint in determining the median CV is that it is dependent on SDSA pairs that occur within a short distance of each other on the same axonal and dendritic segments. This restriction ensures that presynaptic and postsynaptic co-activation results in synapse dimensions that reflect synchronous activation histories (Takahashi et al., 2012). When this constraint is uniformly applied, it becomes possible to determine whether different neural circuits have different information storage capacity in their synapses. We observed previously that the high median CV (∼0.6) of paired spine head volumes in dentate gyrus contrasts dramatically with the much lower median CV (∼0.1) of spine head volumes in hippocampal area CA1 (Bromer et al., 2018; Samavat et al., 2023). This difference may reflect the relatively low basal activity in dentate granule cells compared to CA1 pyramidal cells, and thus a weaker history of co-activation (Bromer et al., 2018), as well as a more hyperpolarized resting membrane potential. Regardless of the source of the differences in CV, when measured in a way that reflects synchronous activation histories, it can be used to determine how much growth (or shrinkage) is required for a particular synapse to achieve a new functionally distinguishable size. Comparing correlations between same-axon same-dendrite, or same-axon same-neuron synapses has been used in large scale cortical connectomics studies to explore structural correlates of functional size (Motta et al., 2019; Dorkenwald et al., 2019). In a connectomics study of adult primary somatosensory cortex, the surface area of the axon-spine interface (ASI) was measured as a proxy for synaptic strength (Motta et al., 2019). When the distribution of CVs among paired synapses were evaluated, large and small spine sizes could be grouped into two categories with equally low CVs (CVs<0.5), whereas random ASI pairs had much higher and more variable CVs (ranging from 0.1-1.5). This outcome led to the conclusion that the SDSA pairs represent potentiated large synapses or depressed small synapses. This finding coincides with our findings of uniform median CV across hippocampal CA1 or dentate granule cell spine head volumes of all sizes. In the primary visual cortex of the adult mouse, outcomes suggested that spine head volumes could be distinguished in two broad peaks of small and large synapses (Dorkenwald et al., 2019). Based on this bimodal distribution, it was suggested that synaptic plasticity switches the synaptic strength between two bistable states. However, within each peak of the bimodal distribution, the sizes of paired synapses across the dendritic tree were highly correlated, and only the frequency of pair sizes was bimodal. Furthermore, these axon-neuron pairs included those spanning multiple dendrites, not just near neighbors on the same dendrite, as were the pairs in the hippocampus analyzed here and elsewhere (Bartol et al., 2015; Bromer et al., 2018), and in the somatosensory cortex (Motta et al., 2019). The precision (CV) of the synaptic pairs in the visual cortex was not assessed but is likely to represent greater than one bit. Surely there will be differences between synapses in the cortex and the hippocampus, as we have shown between different regions of the hippocampus. Regardless of the sources of differences between these studies, it will be interesting to explore the connectomics synaptic data using our new approach based on the precision of synchronous activation for closely spaced SDSA pairs and Shannon information theory.

In adult rat hippocampal area CA1, small spine outgrowth is stalled while synapses on the largest stable spines are further enlarged during LTP (Bell et al., 2014). This LTP-related finding mirrors findings from live imaging, where small spines with and without synapses are more transient than larger stable spines, all of which have synapses (Zuo et al., 2005; Holtmaat et al., 2009; Berry and Nedevi, 2017). One novel but related finding regarding small spine transience in the dentate gyrus is that the initial small spine elevation at 30 min was reduced by 2 hr after the induction of LTP, a time when the number of large spines increased. While this shift likely reflects strengthening of more synapses at 2 hr than at 30 min, the ultimate effect is a reduction in synaptic information storage capacity as more spines enlarge and reduce the range of available spine sizes.

At present, all of the measures of synaptic size have been used as correlative proxies of synaptic efficacy. Perhaps a better measure would be the size of the presynaptic active zone associated with stabilized AMPARs in postsynaptic nanodomains (Biederer et al., 2017; Jung et al., 2021; Haas et al., 2020). As tissue preparation and imaging approaches improve, especially to increase resolution in z, these and other measures of synaptic strength may reveal an even lower CV in SDSA pairs, and thus greater precision of synaptic plasticity. Other features may also need to be added to identify structural indicators of synaptic strength as well as ongoing plasticity in the pairs. These indicators may include nascent zones that have a postsynaptic density but no presynaptic vesicles, mitochondria, SER, and ribosomes, as these features have been found in the past to occur primarily in larger spines during LTP (Bell et al., 2014; Chrillo et al., 2019). It will be interesting to learn whether the variance among the CVs across pairs actually reflect these indicators of plasticity, such that consolidated bistable synapses can be readily distinguished from those in the process of undergoing graduated plasticity. Such refinement in the structural measures of synaptic strength will help to determine which distributions of distinguishable sizes produce the most efficient use of the Shannon entropy distribution and provide the greatest platform for subsequent plasticity.

In conclusion, new and more rigorous analyses enhanced prior findings at 30 minutes (Bromer et al., 2018) and extended them to 2 hr post-LTP induction. We show that the induction of LTP causes a remarkable and durable increase in the information storage capacity of synapses in the dendritic target zone of the stimulated input on granule cells in the dentate gyrus of awake animals. Future studies should investigate whether this increase in information storage capacity persists as long as the LTP effect itself, which in this circuit can last for many months and up to a year in vivo (Abraham et al, 2002).

## Methods and Materials

### Surgery and electrophysiology

The 30 min dataset was from two male Long-Evans rats aged 121 and 179 days, and the 2 hr dataset was from two male Long-Evans rats aged 150 and 170 days at the time of LTP induction and subsequent perfusion. The animals were surgically implanted using stereotaxic coordinates, as previously described in Bowden et al., 2012. Three wire stimulating electrodes were implanted separately into the medial and lateral perforant pathways in the angular bundle of the LTP hemisphere, and in the medial perforant pathway of the control hemisphere. Recording electrodes were implanted bilaterally in the dentate hilus to obtain field excitatory postsynaptic potentials (fEPSP) in response to stimulation at the three electrodes so as to distinguish medial from lateral perforant pathway inputs. Two weeks after surgery, baseline recording sessions commenced, with animals being in a quiet alert size during their dark cycle. Test pulse stimuli were administered to each pathway as constant-current biphasic square-wave pulses (150 µs half-wave duration) at a rate of 1/30 sec alternating between the three stimulating electrodes. The test pulse stimulation intensity was set to evoke medial path waveforms with fEPSP slopes > 3.5 mV/ms in association with population spike amplitudes between 2 and 4 mV, at a stimulation current ≤ 500 µA. Paired-pulse and convergence tests were used to confirm successful stimulating placements in the medial and lateral perforant pathways (Bowden et al., 2012). On the day of LTP induction, all animals received 30 min of baseline test pulses delivered to the two ipsilateral pathways as well as the contralateral medial perforant pathway. The LTP-inducing delta-burst stimulation protocol was delivered to the ipsilateral medial perforant pathway and consisted of five trains of 10 pulses (250 µs half-wave duration at the same pulse amplitude as for the test pulses) delivered at 400 Hz at a 1 Hz interburst frequency, repeated 10 times at 1 min intervals (Bowden et al., 2012). Test pulse stimulation then resumed in the ipsilateral LTP hemisphere and control responses were monitored in the contralateral hippocampus. For the purposes of this paper, the initial slope of the medial path fEPSP in each hemisphere was measured for each waveform and expressed as a percentage of the average response during the last 15 min of recording before DBS.

### Perfusion and fixation

The animals were perfused at 30 min and 2 hr after the onset of delta-burst stimulation under halothane anesthesia and tracheal supply of oxygen (Kuwajima et al., 2013). The perfusion involved a brief (∼20 s) wash with oxygenated Krebs-Ringer Carbicarb buffer [concentration (in mM): 2.0 CaCl_2_, 11.0 D-glucose, 4.7 KCl, 4.0 MgSO_4_, 118 NaCl, 12.5 Na_2_CO_3_, 12.5 NaHCO_3_; pH 7.4; osmolality, 300–330 mmol/kg], followed by fixative containing 2.0% formaldehyde, 2.5% glutaraldehyde (both aldehydes from Ladd Research), 2 mM CaCl_2_, and 4 mM MgSO_4_ in 0.1 M cacodylate buffer (pH 7.4) for ∼1 hr (∼1,900 mL of fixative was used per animal). The brains were removed from the skull at about 1 hr after end of perfusion, wrapped in several layers of cotton gauze, and shipped on ice in the same fixative from the Abraham laboratory in Dunedin, New Zealand, to the Harris laboratory in Austin, Texas by overnight delivery (TNT Holdings B.V.).

### Tissue processing and serial sectioning

The fixed tissue was then cut into parasagittal slices (70 µm thickness) with a vibrating blade microtome (Leica Microsystems) and processed for electron microscopy, as previously described (Kuwajima et al., 2013; Harris et al., 2006). Briefly, the tissue was treated with reduced osmium (1% osmium tetroxide and 1.5% potassium ferrocyanide in 0.1 M cacodylate buffer) followed by microwave-assisted incubation in 1% osmium tetroxide under vacuum. Then the tissue underwent microwave-assisted dehydration and *en bloc* staining with 1% uranyl acetate in ascending concentrations of ethanol. The tissue was embedded into LX-112 epoxy resin (Ladd Research) at 60°C for 48 hr before being cut into series of ultrathin sections at the nominal thickness of 45 nm with a 35° diamond knife (DiATOME) on an ultramicrotome (Leica Microsystems). The serial ultrathin sections were obtained in the middle of the medial perforant pathway input located ∼125 µm from the top of the granule cell layer in the dorsal blade of the hippocampal dentate gyrus. The sections were collected onto Synaptek Be-Cu slot grids (Electron Microscopy Sciences or Ted Pella) that were coated with Pioloform (Ted Pella) and stained with a saturated aqueous solution of uranyl acetate followed by lead citrate (Reynolds, 1963).

### Imaging and alignment

The serial ultrathin sections were imaged, blind as to condition, with either a JEOL JEM-1230 TEM or a transmission-mode scanning EM (tSEM) on the Zeiss SUPRA 40 field-emission SEM with a retractable multimode transmitted electron detector and ATLAS package for large-field image acquisition (Kuwajima et al., 2013). On the TEM, sections were imaged in two-field mosaics at 5,000 times magnification with a Gatan UltraScan 4000 CCD camera (4,080 pixels × 4,080 pixels), controlled by DigitalMicrograph software (Gatan). Mosaics were then stitched with the photomerge function in Adobe Photoshop. On the tSEM, each section was imaged with the transmitted electron detector from a single field encompassing 32.768 µm × 32.768 µm (16,384 pixels × 16,384 pixels at 2 nm/pixel resolution). The scan beam was set for a dwell time of 1.25–1.4 µs, with the accelerating voltage of 28 kV in high-current mode. The TEM mosaics and tSEM images were aligned through serial sections using Fiji with the TrakEM2 plugin (Cardona A, et al., 2012; Schindelin J, et al., 2012); fiji.sc). The images were aligned rigidly first, followed by application of affine and then elastic alignment as needed.

### Unbiased sampling of dendritic segments and associated dendritic spines

Images from a series were given a five letter code to mask the identity of experimental conditions in subsequent analyses with the Reconstruct software (Fiala, 2005). Pixel size was calibrated for each series using the grating replica image that was acquired along with serial sections. The section thickness was estimated using the cylindrical diameters method (Fiala et al., 2001). All segmentations and curations were done by experienced tracers who remained blind as to the experimental conditions until the codes were revealed for statistical and graphical analyses. Three dendrites were traced through serial sections from each control and LTP hemisphere at each time point for a total of six dendrites per condition with a total of 24 dendrites over all conditions. Dendritic spine density has previously been shown to scale with dendrite cross-sectional area and microtubule count (Harris et. al., 2022). Hence, dendrites were randomly chosen from the population that spanned the image volume and had a microtubule count ranging from 30 to 35, which was within 2 counts from the average count among all dendrites in the middle molecular layer of the dentate gyrus (Harris et. al., 2022). The z-trace tool in Reconstruct was used to obtain the unbiased lengths spanning the origin of the first included spine to the origin of the last spine (Fiala et al., 2001). These dendritic segments ranged in length from 8-11 µm across all series and were similar for all conditions (Table 1).

### Generating spine head volumes

Spine head volumes were chosen as a proxy for synaptic strength because current 3D methods provide more reliable measurements than other correlated metrics, such as synapse area or vesicle number (Bartol et al., 2015). Dendritic spines were segmented as previously described using the Neuropil Tools analyzer tool in Blender, a free, open-source, user-extensible computer graphics tool in conjunction with the 3D models generated in Reconstruct (Bartol et al., 2015). Filtered selection of traced objects from Reconstruct files were used to generate 3D representations of selected contours in Blender by invoking functions from VolRoverN (Edwards et al., 2014). Thus, mesh objects were generated with triangle faces between contour traces. Smoothing the surface of the spines was accomplished with GAMer software fetk.org/codes/gamer/). In a few cases, the formation of triangles was uneven and required additional manipulation by Blender tools and then repeating this smoothing process. For purposes of visualization, the synaptic locations are represented as patches of red colored triangles and tagged with identification metadata in the reconstructed surface mesh of the spine heads. The selection of spine head from spine neck was made using the 3D visualization in Blender to identify the halfway point along the concave arc as the head narrowed to form the neck (Bartol et al., 2015; Bromer et al., 2018). To ensure the accuracy of the measurements, segmentation of each spine head volume was performed four times (twice each by two people) and the average and CV of the four measurements was calculated. A further check was added when spine heads with a CV ≥ 0.02 for the four measurements were visually evaluated by an expert and any error in the segmentation was corrected.

### Identification of same-dendrite same-axon (SDSA) Pairs

The trajectories of the presynaptic axons are such that they do not bend back and forth but instead pass by the dendrites at an angle and sometimes form synapses with more than one spine on the same dendrite (SDSA pairs). Only spines located in the middle of the sample dendrite had presynaptic axons sufficiently complete within the series to determine connectivity. The axons were traced past the nearest neighboring axonal boutons on either side of the one that synapses with a central spine, until it was determined whether they formed synapses with the same or different dendrites. At least 15 axons were evaluated per dendrite. In some cases, the spines were branched but counted as one spine origin with multiple presynaptic axons that were evaluated separately in the SDSA pairs analysis. Some spines were multisynaptic with either two excitatory (asymmetric) synapses or one excitatory and one inhibitory (symmetric) synapse. Only the axons forming excitatory synapses were included in this SDSA pairs analysis. Occasionally, excitatory, or inhibitory shaft synapses were interspersed along the middle segment and these were also excluded in the SDSA pairs analysis since shaft synapses have no spine head volumes. If a SDSA paired spine was discovered beyond the central analysis of 15 axons, both it and its partner were included in the SDSA pairs analysis and the analyzed dendritic length was extended to include the intermediate spines. Three axons made synapses with dendritic spines of two different sample dendrites and they were all included for frequency calculations. This central dataset was used to evaluate SDSA pairs across conditions (Table 1).

### Reanalysis of published data

The 30 min dataset is fully described above and was also analyzed in a prior paper using signal detection theory (Bromer et al., 2018), an approach that is not as robust as the new approach based on Shannon information theory (Samavat et al., 2023). (Note: Earlier versions of this new method appeared in (***Samavat et al., 2022; Samavat et al., 2022; Samavat et al., 2022; Samavat M, 2023***)).

## Statistical Analysis

### Precision analysis of SDSA pairs

Precision was defined as the degree of reproducibility of a measurement (Samavat et al., 2023). The CV shown in equation 1 is a statistic that measures variations within a sample. Standard deviation (σ) was defined in equation 2 and the CV was normalized by the mean of the sample (μ). Here *N* = 2 in equation 2 for each SDSA pair.

### Standard error bootstrapping of the median for SDSA pairs

The standard error of median for the precision levels of each of the SDSA pairs in 4 datasets was calculated by generating 10,000 bootstrap samples of size n, each sampled from the n SDSA pairs with replacement, to estimate the standard error of the median for the n SDSA pairs (gray boundaries in Fig. 3, and column 3 of Table 1). Computing the standard error of median of spine head volumes followed the same procedure (For details see Algorithm 1 in Samavat et al., 2023). The standard errors of the entropy, efficiency constant, maximum entropy for uniform distribution, and KL divergence (Table 2 column 2-5) were all calculated using this bootstrapping technique. Also, see Efron et al., 2021 for additional justification of bootstrapping for the calculation of standard error.

### Distribution of distinguishable sizes based on spine head volumes

To construct the distribution of distinguishable sizes, spine head volumes were first sorted from smallest to the largest. The first value (smallest value) was selected and the CV of that value and the remaining head volumes were calculated in a pairwise manner. The head volumes for which the calculated CV was below the threshold (the median value of the SDSA pairs CV) were assigned to the first size and deleted from the pool of N spine head volumes. This procedure was repeated until the CV exceeded the median SDSA pairs CV and a new size bin was then formed. New sizes were formed until all the remaining spine head volumes were assigned and the original vector of spine head volumes was empty (Algorithm 2, Samavat et al., 2023). The coefficient of variation between each random pair within each size was less than the threshold value measured from the reconstructed SDSA pairs.

### Kullback-Liebler (KL) divergence and efficiency ratio

Measurement of the distance between an observed distribution with a reference probability distribution can be done by Kullback-Liebler (KL) divergence. A uniform distribution is the discrete probability distribution with maximum entropy when there is no constraint on the distribution except having the sum of the probabilities equal 1 for a fixed number of sizes. Formally, the KL divergence between the distribution of spine head volume sizes (P) and the uniform distribution of sizes (U) is the difference between cross entropy of (P) and (U) and the entropy of (P): [*H*(*P*,U) − *H*(*P*)]. The closeness of fit between the distribution of the distinguishable sizes to the maximum entropy distribution was determined with the fixed number of sizes as the only known constraint on the distribution. When the distribution of the distinguishable synaptic sizes approaches the uniform distribution, the KL divergence decreases and the Shannon entropy will approach maximum.

The outcomes provide an upper bound for the sampled synapses but not necessarily for other synapses. We also calculated the ratio of the KL divergence values over the maximum value that KL divergence can possibly have (KL / KL_MAX_) where KL_MAX_ equals the entropy of a uniform distribution, *H*(*U*). This ratio was then used to define the efficiency ratio η=(1-(KL / KL_MAX_))*100. It measures the efficiency of the measured KL divergence value with respect to the minimum value it could hypothetically have. The higher the efficiency ratio, the more efficient is the usage of distinguishable sizes for the storage of synaptic strength values across the population of synapses.

### General statistical approaches

Statistical analysis and plots were generated using Python 3.4 with NumPy (Harris et al., 2020), SciPy (Virtanen et al., 2020), and Matplotlib (Hunter, 2007). R package ggplot2 (Wickham, 2011), ggpubr, scales, xlsx, and ggpmisc. The coefficient of determination, denoted R^2^, was used in Fig. 3 to show the proportion of the variation in the dependent variable (CV) that is predictable from the independent variable (spine head volumes). Lognormal transformation of data was performed on skewed distributions (Fig. 2 A-F, Fig. 5 A-D).

## Acknowledgments

This research is part of a multi-institutional collaboration project funded by the NSF NeuroNex Technology Hub (DBI-1707356) and Next Generation Networks for Neuroscience (DBI-2014862) led by KMH. This research was also supported by NSF IIS-2219979; NIH P41GM103712; NIH MH095980; NIH MH115556; NIH MH129066.

## Author contributions

MS, TMB, WCA, KMH, and TJS designed research; MS, TMB, WCA, KMH, and TJS analyzed data; MS designed and implemented all the simulation algorithms and generated results with input from TMB, WCA, KMH, and TJS. WCA,JBB, KMH, MK, and JM designed and performed electrophysiology experiments, tissue processing, and imaging. MS, DDH, DCH, MK, PHP, and KMH performed and curated reconstructions and prepared materials for Fig. 1. MS, TMB, KMH, WCA, and TJS wrote the paper with input from all authors. For more details about the physiology, WCA is available for correspondence at.

## Data and Code Availability

The codes for algorithms used in the present study will be available in the following github link upon publication: https://github.com/MohammadSamavat. Data will be made publicly available upon publication via 3dem.org (DOI will be added at time of publication).

---

**Supplemental Table 1:** Metadata and analyses for each time point, animal, and condition.

| *time points* | 30 min |  |  |  | 2 hours |  |  |  |
| --- | --- | --- | --- | --- | --- | --- | --- | --- |
| *animal # (ID)* | Animal 1 (LED50) |  | Animal 2 (LED56) |  | Animal 3 (LED108) |  | Animal 4 (LED113) |  |
| *weight (gr)* | 623 |  | 448 |  | 648 |  | 490 |  |
| *age (days)* | 179 |  | 121 |  | 170 |  | 150 |  |
| *circadian cycle* | dark |  | dark |  | dark |  | dark |  |
| *time of day* | 10:30 am |  | 10:30 am |  | 09:30 am |  | 09:30 am |  |
| *series code* | CLZBJ | TNKPS | NDKZB | MFBCF | JHHZS | HLWLQ | KSGRS | BBCHZ |
| *condition* | control | LTP | control | LTP | control | LTP | control | LTP |
| *ave pre % fEPSP* | -3.84 | -3.81 | 3.16 | 2.81 | -2.36 | 0.04 | 0.65 | -2.53 |
| *ave post % fEPSP* | -5.38 | 33.61 | 5.28 | 48.33 | -4.76 | 40.68 | -1.10 | 34.45 |
| *# SDSA pairs* | 6 | 4 | 4 | 4 | 10 | 4 | 8 | 6 |
| *median CV* | 0.65 | 0.31 | 0.71 | 0.76 | 0.44 | 0.42 | 0.63 | 0.38 |
| *# spines* | 130 | 110 | 79 | 78 | 112 | 111 | 127 | 115 |
| *# distinguishable synaptic strength* | 5 | 11 | 4 | 5 | 6 | 8 | 5 | 9 |
| *#bits* | 1.97 | 3.1 | 1.77 | 1.93 | 2.16 | 2.68 | 1.92 | 2.89 |
| *KL div.* | 0.35 | 0.36 | 0.23 | 0.40 | 0.43 | 0.32 | 0.41 | 0.28 |
| *# bootstrap distinguishable strength* | 4±0.47 | 10±0.55 | 4±0.39 | 4±0.5 | 6±0.38 | 7±0.71 | 5±0.38 | 8±0.6 |
| *# bootstrap bits* | 1.92±0.06 | 2.98±0.1 | 1.7±0.13 | 1.86±0.1 | 2.1±0.1 | 2.54±0.1 | 1.86±0.1 | 2.77±0.1 |
| *bootstrap KL div.* | 0.14±0.12 | 0.31±0.09 | 0.24±0.12 | 0.27±0.13 | 0.46±0.1 | 0.27±0.1 | 0.42±0.12 | 0.31±0.08 |

The pre % fEPSP was averaged over the five minutes prior to DBS, and the post % fEPSP was averaged over the last five minutes prior to perfusion fixation. KL div. is KL divergence. The median±SE in the bottom three rows was calculated with bootstrapping. All animals were on a reverse light/dark circadian cycle.

